# A quantitative metric of pioneer activity reveals that HNF4A has stronger in vivo pioneer activity than FOXA1

**DOI:** 10.1101/2022.05.24.493302

**Authors:** Jeffrey L Hansen, Barak A Cohen

## Abstract

We and others have suggested that pioneer activity–a transcription factor’s (TF’s) ability to bind and open inaccessible loci–is not a qualitative trait limited to a select class of pioneer TFs. We hypothesize that most TFs display pioneering activity that depends on the TF concentration and the motif content at their target loci. Here we present a quantitative measure of pioneer activity that captures the relative difference in a TF’s ability to bind accessible versus inaccessible DNA. The metric is based on experiments that use CUT&Tag to measure binding of doxycycline (dox) inducible TFs. For each location across the genome we determine a “dox_50_,” the concentration of dox required for a TF to reach half-maximal occupancy. We propose that the ratio of a TF’s average dox_50_ between ATAC-seq labeled inaccessible and accessible binding sites, its Δdox_50_, is a measure of its pioneer activity. We measured Δdox_50_’s for the endodermal TFs FOXA1 and HNF4A and show that HNF4A has a smaller Δdox_50_ than FOXA1, suggesting that HNF4A has stronger pioneer activity than FOXA1. We further show that FOXA1 binding sites with more copies of its motif have a lower Δdox_50_, suggesting that strong motif content may compensate for weak pioneer activity. The quantitative analysis of binding suggests different modes of binding for FOXA1, including an anti-cooperative mode of binding at certain accessible loci. Our results suggest that Δdox_50_s, or other similar measures that assess the difference in TF affinity for inaccessible and accessible DNA, are reasonable measures of pioneer activity.

## Introduction

Activating silent genes requires transcription factors (TFs) to bind and open DNA when their motifs are occluded by nucleosomes. Activating silent genes is postulated to involve two qualitatively different classes of TFs, pioneer factors (PFs) and non-pioneer factors (nonPFs) (Cirillo et al. 2002; Iwafuchi-Doi and Zaret 2014). According to this hypothesis PFs bind to nucleosome-occluded DNA and make it accessible to nonPFs, which then recruit the cofactors required to activate transcription. However, we recently showed that both a canonical PF, FOXA1, and a nonPF, HNF4A, can independently bind, open, and then activate nearby genes (Hansen, Loell, and Cohen 2022), and many TFs possess unique ways of binding and opening nucleosomal DNA (Zhu et al. 2018; Swinstead et al. 2016; Miller and Widom 2003; Soufi et al. 2015; Yu and Buck 2019). From these data we propose that most TFs have quantifiable pioneer activity that depends on their nuclear concentrations and the motif content at their target loci. Here we present a metric that quantifies the pioneer activity of TFs at loci across the genome. The distribution of pioneer activity across the genome supports the hypothesis that most TFs can pioneer silent loci given appropriate TF levels and motif content.

## Results

### Definition of “Δdox_50_” parameter for pioneer activity

An appropriate measure of pioneer activity should capture the relative difference of TF binding between inaccessible and accessible sites in the genome. In principle, we could compare the dissociation constant (K_d_) of a TF at inaccessible and accessible sites as a measure of pioneer activity, since the K_d_ is the concentration of TF required to reach half maximal binding. In practice, computing a K_d_ inside cells is impractical because it requires measuring the absolute concentration of a TF in the nucleus, in its proper post-translationally modified state. We propose a related measure that uses doxycycline-inducible (dox-inducible) TFs to compute the dox_50_, the dox concentration required to reach half-maximal binding inside cells. By inducing TF levels over a wide range of dox concentrations and measuring the resulting binding by CUT&Tag (Kaya-Okur et al. 2019), we determine a dox_50_ for every location in the genome in parallel. The ratio of the average dox_50_ at inaccessible versus accessible sites, its Δdox_50_, is a quantitative measure of a TF’s pioneering activity. The smaller a TF’s Δdox_50_, the less its binding is reduced at inaccessible DNA (Fig. 1A). Because the measurements at inaccessible and accessible sites are made at the same time in the same nucleus, the dox concentrations (or TF concentrations) cancel out, allowing us to compare the Δdox_50_s of different TFs to each other (Man and Stormo 2001). This strategy allows us to circumvent the challenge of measuring effective nuclear TF concentration while maintaining the physiological relevance of our *in vivo* pioneer activity measurements.

**Figure 1.**
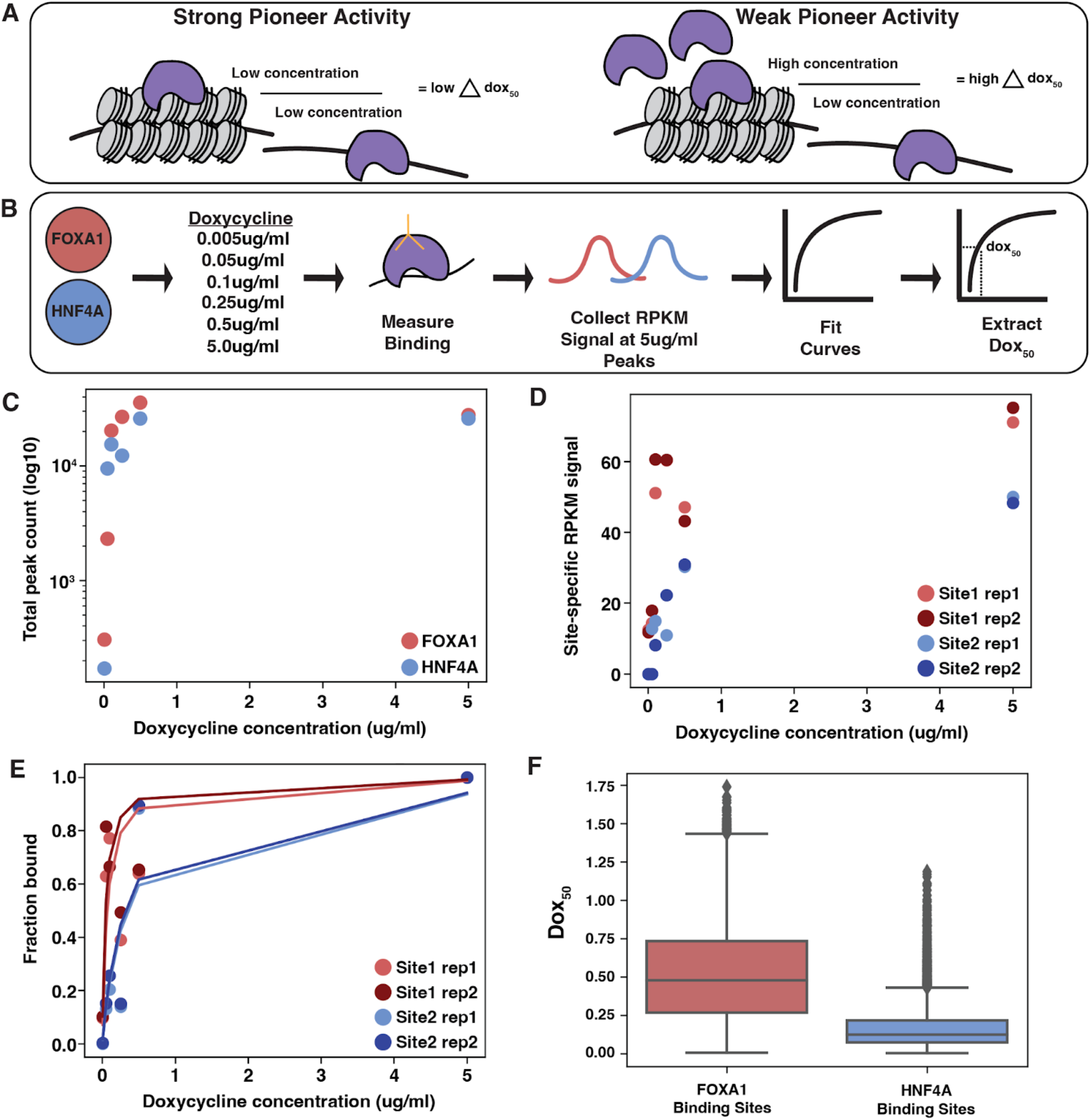
Experimental design to calculate TF dox_50_ values across the genome. **(A)** Strong pioneer activity leads to smaller differences in the dox concentration needed to achieve binding at inaccessible versus accessible sites, and vice versa. **(B)** We induced FOXA1 or HNF4A across a 1,000-fold dox range, measured binding, collected binding signal at a reproducible set of binding sites and then extracted a dox_50_ for each site. **(C)** Number of total peaks for each TF across each dox induction level. **(D)** Replicate RPKM binding signal at two example genomic sites. **(E)** Replicate fitted lines at two example genomic sites. **(F)** Full distribution of dox_50_ values for each TF.

### Measurement of dox_50_ for FOXA1 and HNF4A

FOXA1 and HNF4A are liver TFs that are commonly used to reprogram embryonic fibroblasts to endoderm progenitor cells (Biddy et al. 2018; Sekiya and Suzuki 2011). FOXA1 is a canonical PF and HNF4A a nonPF, and the two are suggested to work in a collaborative and sequential fashion to activate their target genes (Horisawa et al. 2020; Cirillo et al. 2002). We previously tested FOXA1 and HNF4A’s behavior in an ectopic setting by expressing them within K562 blood cells, a lineage in which neither TF is expressed and that should present the TFs with unique complements of chromatin and cofactors. We created clonal K562 lines that expressed either inducible FOXA1 or HNF4A and showed that both TFs could independently bind and open inaccessible chromatin and activate nearby genes (Hansen, Loell, and Cohen 2022).

Based on the ability of FOXA1 and HNF4A to independently bind, open, and activate in an ectopic cell line, we expected both TFs would have similar pioneer activity. To test this prediction we attempted to measure each TF’s Δdox_50_ using the same dox-inducible FOXA1 or HNF4A K562 lines (Fig. 1B). We first treated each TF line with a 1,000-fold range of dox (0.005, 0.05, 0.25, 0.1, 0.5, and 5.0μg/ml) and measured resultant binding. Read normalized binding signal (RPKM) was highly correlated between replicates (Fig. S1). FOXA1 and HNF4A appear to be expressed at similar levels at each dox level as each TF bound to similar numbers of sites in each condition (Fig. 1C). We then collected the overlapping set of binding sites between each TF’s replicates in the 5.0μg/ml sample and at each site plotted the read normalized signal (RPKM) from the other induction levels (Fig. 1D). Generally the binding patterns follow predicted saturation binding kinetics (Fig. S2). We then fit Equation 1 to these distributions.

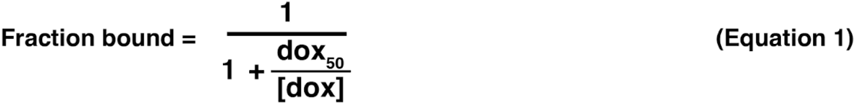

In order to fit Equation 1, we normalized each site’s RPKM signal to the signal in the 5.0μg/ml sample to convert our measurements into fractional binding (Fig. 1E). We found that at some sites the binding signal peaked prior to the 5.0μg/ml sample (fraction bound > 1 in any of the first five induction levels). We removed these sites to prevent poor fitting. This left us with 11,557 FOXA1 binding curves and 5,940 HNF4A binding curves with highly similar fitted lines across replicates (Fig. 1F, Fig. S3). We extracted dox_50_ values from these lines and found similar results between replicates (Fig. S4) and so we averaged each site’s replicate dox_50_ value for the remaining analyses. The resulting distributions of each TF’s genome-wide dox_50_ values show that FOXA1 has a much larger variance in dox_50_ values than HNF4A (Fig. 1F), suggesting that FOXA1 binding generally depends more on the genomic environment than HNF4A.

### Measurement of Δdox_50_ for FOXA1 and HNF4A

The dox_50_ distributions in Figure 1 suggest that HNF4A may bind more consistently across the genome but do not explicitly measure their pioneer activities, the difference in each TFs ability to bind at inaccessible versus accessible sites. We therefore classified each site as either inaccessible or accessible based on ATAC-seq (Buenrostro et al. 2015) peaks collected in these cell lines before dox induction (Hansen, Loell, and Cohen 2022). Of FOXA1’s 11,557 peaks, 1,930 were in accessible regions and 9,627 were in inaccessible regions (10,120 accessible before filtering, 17,644 inaccessible before filtering). Of HNF4A’s 5,940 peaks, 2,135 were in accessible regions and 3,805 were in inaccessible regions (16,137 accessible before filtering, 16,507 after filtering). Comparing the dox_50_ distributions between inaccessible and accessible sites revealed that the binding of HNF4A is less affected by inaccessible DNA than FOXA1 (Fig. 2A). We then computed a Δdox_50_ for each TF by dividing the average dox_50_ for inaccessible sites by the average dox_50_ for accessible sites. This analysis showed that HNF4A has a lower Δdox_50_ than FOXA1 (Table 1). This result holds when we include those sites that were filtered out because they peaked at lower concentrations (Table 1, Fig. S5).

**Figure 2.**
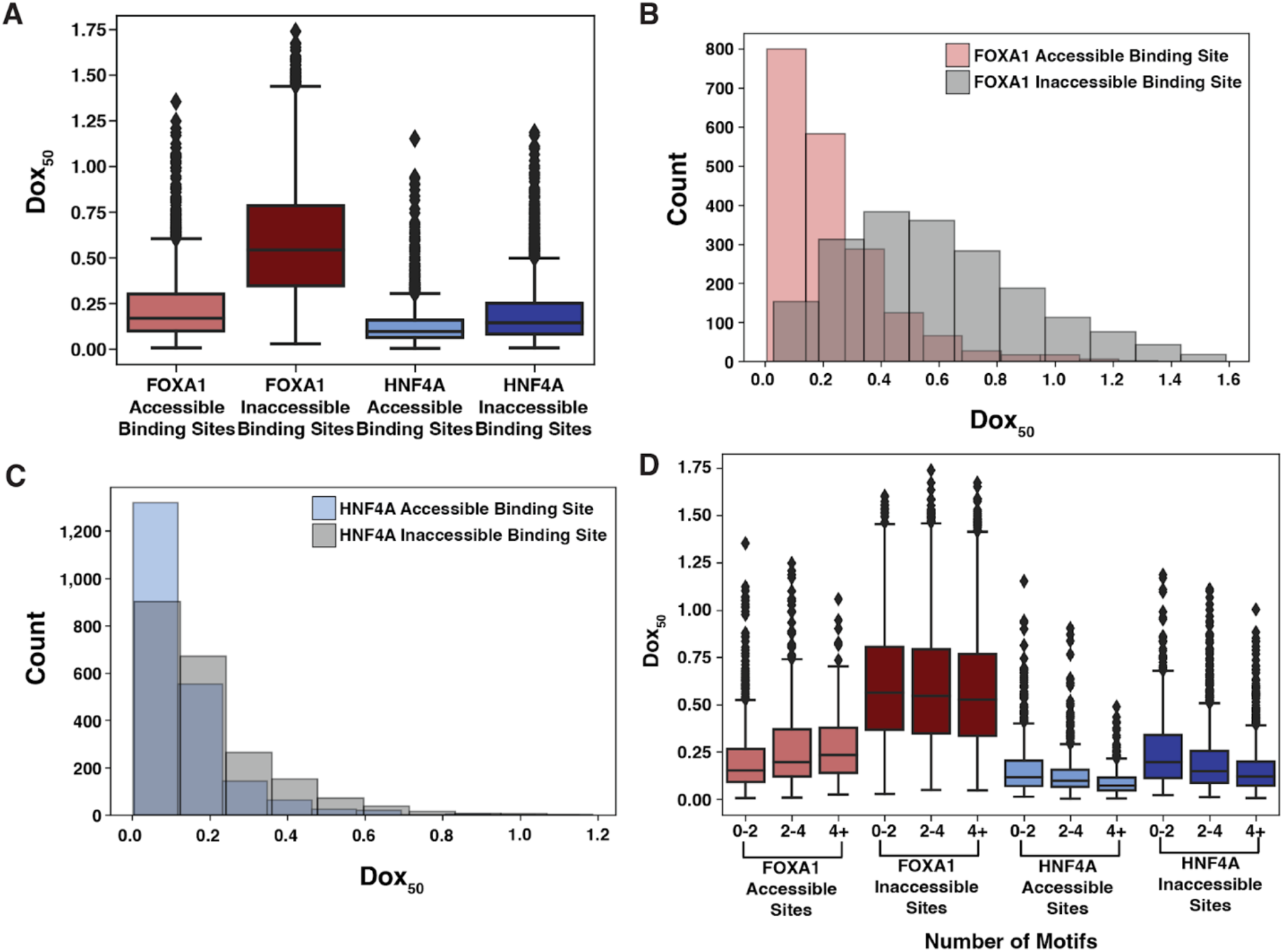
HNF4A has a smaller Δdox_50_ than FOXA1. **(A)** Distributions of dox_50_ estimates extracted from binding curves at FOXA1 accessible binding sites (*n* = 1,930), FOXA1 inaccessible binding sites (*n* = 9,627), HNF4A accessible binding sites (*n* = 2,135), and HNF4A inaccessible binding sites (*n* = 3,805). **(B-C)** Distributions from FOXA1 **(B)** and HNF4A **(C)** shown in histogram form. **(D)** Same plot as (A) but each genomic binding site is binned by whether the site has < 2, >= 2 but < 4, or >= 4 motifs as called by FIMO (*p* = 1e-3).

**Table 1.** Δdox_50_s for FOXA1 and HNF4A across different types of binding sites.

| TF | All sites | Filtered sites | $< 2$ motifs | 2-4 motifs | $> 4$ motifs |
| --- | --- | --- | --- | --- | --- |
| FOXA1 | 5.23 | 2.56 | 2.96 | 2.16 | 2.02 |
| HNF4A | 1.81 | 1.47 | 1.55 | 1.56 | 1.70 |

We next considered whether the motif content at each binding site affected the Δdox_50_. We subset each TF’s binding sites into those that had less than 2, between 2-4, or more than 4 copies of its respective motif and re-plotted the dox_50_ distributions and re-calculated Δdox_50_s. Higher motif content correlated with more consistent binding of FOXA1 between inaccessible and accessible sites and thus lowered FOXA1’s Δdox_50_ (Fig. 2D, Table 1). In contrast, the number of HNF4A motifs at a target locus had little effect on HNF4A (Fig. 2D, Table 1). We conclude that HNF4A has stronger pioneer activity in K562 cells and that the weaker pioneer activity of FOXA1 can be compensated by strong motif content.

### Chromatin modifications explain some of the variance in dox_50_ values

We built a linear model (Equation 2) to try to explain the variance in dox_50_s for FOXA1 and HNF4A where C(Accessibility) is each binding site’s accessibility prior to TF induction. Accessibility explained 17% of the variance in FOXA1’s dox_50_s but only 4% of HNF4A’s. While these data further underscore the greater role that accessibility plays on FOXA1 binding than HNF4A, they also reveal that most of the variance in dox_50_ values between genomic loci must be explained by some other variable.

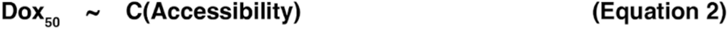

We hypothesized that some of the remaining variance may be explained by the different chromatin modifications present at different target loci and predicted that binding sites with active marks would have lower dox_50_ distributions (easier binding) and binding sites with silent marks would have higher dox_50_ distributions (harder binding). We further subset each TF’s accessible or inaccessible binding sites into those that overlap common K562 marks (Zhang et al. 2020). H3K4me1 marks enhancers (Heintzman et al. 2007), H3K27Ac marks activity (Creyghton et al. 2010), and H3K9me3 and H3K27me3 are two modifications shown previously to suppress pioneer activity (Mayran et al. 2018). The accessible sites overlapped much more often with active marks than silencing marks, and vice versa, and we found that no FOXA1 or HNF4A accessible sites were marked with H3K9me3 (Fig. 3).

**Figure 3.**
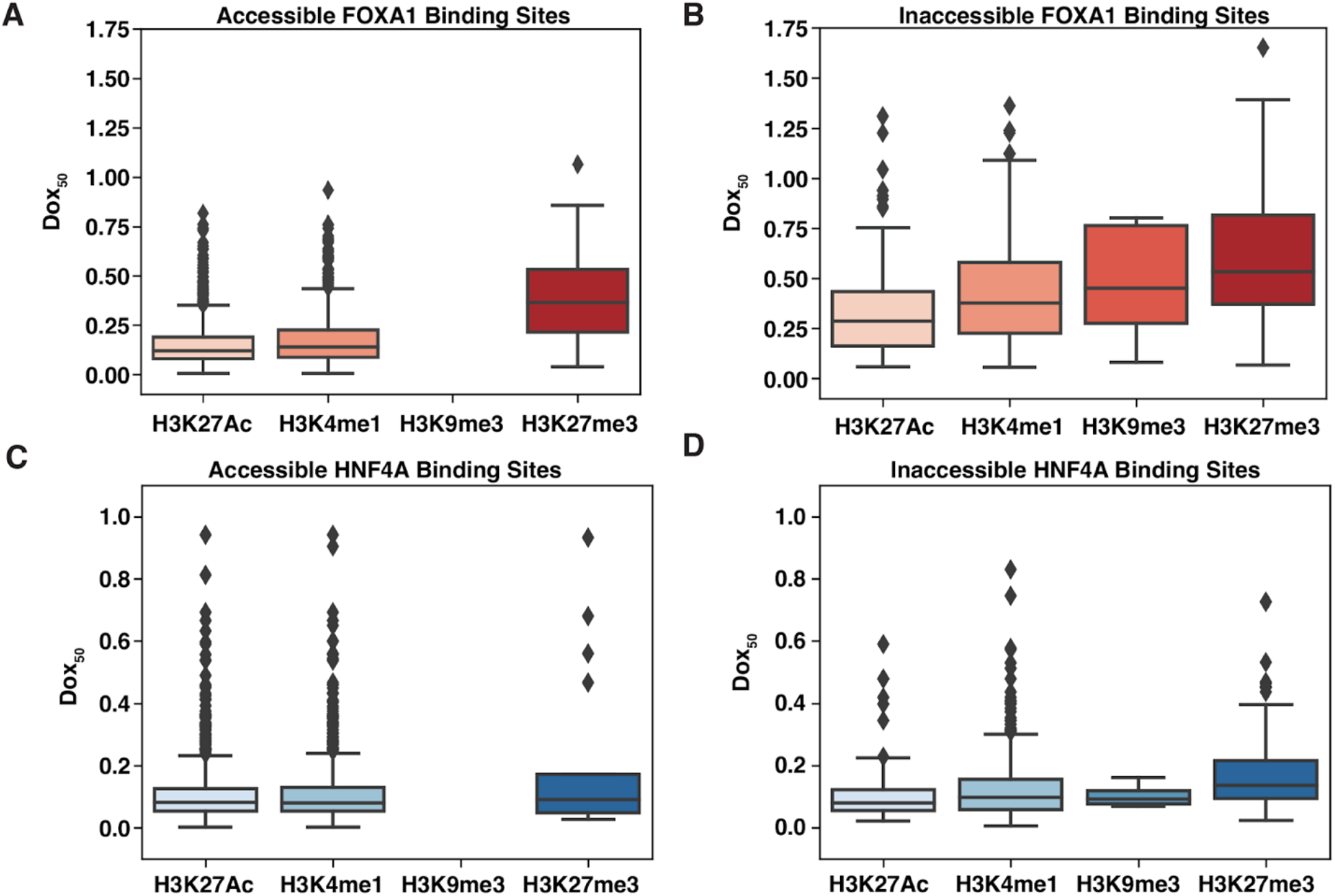
Dox_50_ distributions across different chromatin modifications. **(A)** Dox_50_ values for FOXA1 accessible binding sites that overlapped H3K27AC (*n* = 1,288), H3K4me1 (*n* = 755), H3K9me3 (*n* = 0), and H3K27me3 (*n* = 21). **(B)** Dox_50_ values for FOXA1 inaccessible binding sites that overlapped H3K27AC (*n* = 203), H3K4me1 (*n* = 352), H3K9me3 (*n* = 12), and H3K27me3 (*n* = 277). **(C)** Dox∞ values for HNF4A accessible binding sites that overlapped H3K27AC (*n* = 1,147), H3K4me1 (*n* = 1,111), H3K9me3 (*n* = 0), and H3K27me3 (*n* = 17). **(D)** Dox_50_ values for HNF4A inaccessible binding sites that overlapped H3K27AC (*n* = 140), H3K4me1 (*n* = 416), H3K9me3 (*n* = 4), and H3K27me3 (*n* = 135).

As we predicted, FOXA1 or HNF4A binding sites that overlapped H3K27Ac or H3K4me1 chromatin modifications had lower dox_50_ distributions than those that overlapped H3K9me3 or H3K27me3 (Fig. 3). These effects were present even after we subset binding sites by accessibility, suggesting that the chromatin modifications can affect binding in ways that are separable from the effects of accessibility. However, when we individually added each chromatin modification (plus an interaction term) to the model in Equation 2, we found that accounting for these marks did not have large effects on the ability of the model to predict dox_50_ values for either TF. H3K27ac levels explained 2% of FOXA1’s dox_50_ variance, H3K4me1 explained 1%, and H3K27me3 explained <1%. For HNF4A, H3K27ac explained 2%, H3K4me1 explained 2%, and H3K27me3 explained <1%. All interaction terms were negligible. Together these data suggest that something besides the epigenetic landscape of loci is having a large effect on the pioneering activity of TFs.

### FOXA1 behaves anti-cooperatively at a subset of accessible binding sites

While examining individual binding sites and their fitted curves, we observed a repeating pattern at a subset of genomic locations where the binding signal increased to a peak at the third (0.1μg/ml) or fourth (0.25μg/ml) induction level and then decreased at the highest dox concentration, suggesting anti-cooperative behavior (Fig. 4A, Fig. S6). To quantify the prevalence of anti-cooperative binding, we sampled 10,000 peaks from FOXA1 or HNF4A inaccessible or accessible binding sites and then counted how many displayed saturation behavior (peak at 5μg/ml, Fig. S2) and how many displayed anti-cooperative behavior (peak at 0.1μg/ml or 0.25μg/ml, Fig. S6). We found that the anti-cooperative behavior occurs most often at accessible FOXA1 binding sites (Fig. 4B-C). Anti-cooperative behavior does not appear to depend on the number of motifs at each peak (Fig. 4D) or the length of each peak (Fig. 4E).

**Figure 4.**
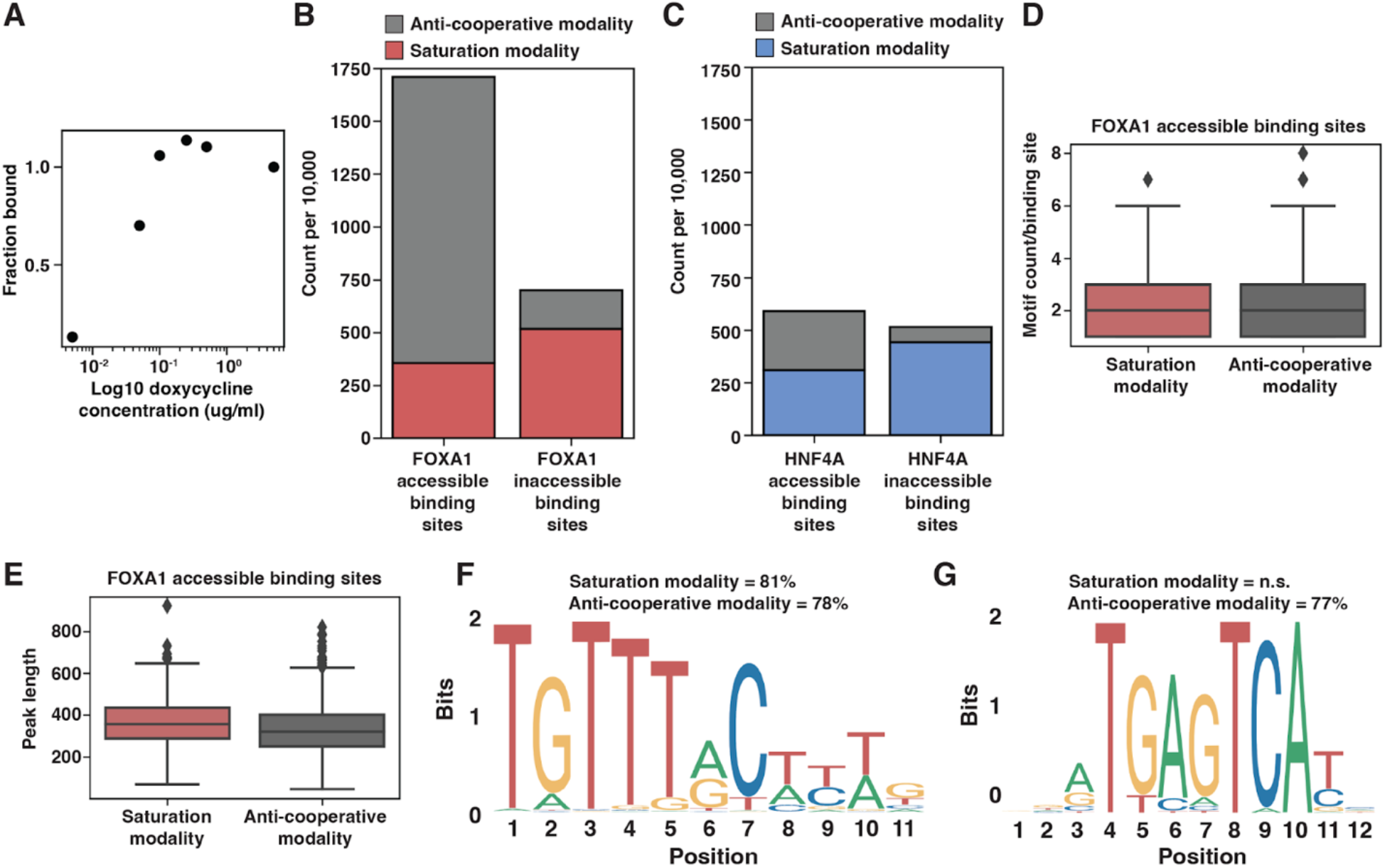
Characterization of anti-cooperative binding behavior. **(A)** Example binding curve at a single genomic site that exhibits anti-cooperative behavior. **(B-C)** A sample of 10,000 FOXA1 **(B)** or HNF4A **(C)** accessible (left bar) or inaccessible (right bar) binding sites colored by if they display saturation binding behavior (gray) or anti-cooperative binding behavior (red, see). **(D)** FOXA1 motif count between the accessible binding sites from (B) that display either saturation or anti-cooperative binding behavior. Motifs were called from FIMO with a p-value threshold of 1e-3. **(E)** Binding peak length between the accessible binding sites from (B) that display either saturation or anti-cooperative binding behavior. **(F)** The most enriched motif discovered in FOXA1 accessible saturation and anti-cooperative peaks was FOXA1 (JASPAR MA0148.1). It is significantly enriched for both the saturation behavior (*p* = 2.19e-001) and anti-cooperative behavior (*p* = 1.23e-01). **(G)** The second most enriched motif discovered in FOXA1 accessible anti-cooperative peaks was AP1 (JASPAR MA1141.1). It was not discovered in the saturation behavior peaks. It is significantly enriched for only the anti-cooperative behavior (*p* = 1e-008).

We considered whether another TF might be contributing to anti-cooperative behavior by searching for enriched motifs in either saturation-type accessible FOXA1 binding sites or anti cooperative-type sites. While FOXA1 sites were enriched in both types of loci (Fig. 4F), the AP1 motif was only enriched at anti-cooperative sites (Fig. 4G). AP1 is an important K562 TF that exhibits some pioneer activity (Biddie et al. 2011). The results suggest that a genetic interaction between FOXA1 and AP1 underlies anti-cooperative behavior at accessible loci.

## Discussion

Given a definition of pioneer activity that is a TF’s ability to bind at inaccessible genomic locations, we suggest that the Δdox_50_ is a quantitative measure of this activity. We measured the Δdox_50_ of FOXA1 and HNF4A in K562 cells and showed that HNF4A has stronger pioneer activity in this cell type than FOXA1. However, both TFs showed a range of dox_50_ values across the genome, which demonstrates that a TFs pioneer activity may depend on accessibility, native chromatin marks, and other factors. Some of these differences are explained by the motif content at different locations, suggesting that low pioneer activity can be overcome by strong motif content. While our work shows that the pioneering activity of a TF can vary across the genome, what accounts for this variation across sites remains mostly unexplained. DNA accessibility had the largest effect on pioneer activity but only explained 17% of the variance in dox_50_ values. We speculate that much of the remaining variance in dox_50_ values might be explained by interactions with other specific TFs or with the general transcription machinery that can differ across the genome.

Our work supports the hypothesis that pioneer activity is not a qualitative trait limited to a few TFs, but rather a quantitative property of TFs that manifests differently depending on the TF and the genomic environment (Garcia et al. 2019; Zhu et al. 2018; Hansen, Loell, and Cohen 2022; Cirillo et al. 2002; Soufi et al. 2015; Yu and Buck 2019). Pioneer activity as a quantitative trait fits with data showing that TFs can gain pioneer activity when expressed at high levels or when cells are forced into replication (Yan, Chen, and Bai 2018) and that TFs can lose pioneer activity when expressed at lower levels (Hansen, Loell, and Cohen 2022).

*In vitro,* FOXA1 has higher affinity (a lower K_d_) for naked DNA than HNF4A (Jiang, Lee, and Sladek 1997; Garcia et al. 2019; Rufibach et al. 2006), and yet HNF4A has stronger pioneer activity (a lower Δdox_50_) than FOXA1 in K562 cells. These results demonstrate that pioneer activity is not solely a function of the affinity of a TF’s DNA-binding domain for its cognate motif. Inside cells, pioneer activity likely depends on the interactions a TF makes with other TFs and with cofactors. Because these interactions will differ in different cell types, a TF’s pioneering activity is also likely to depend on the cell type in which it is expressed and the co-bound TFs present at certain locations.

At some locations in the genome, an interaction between FOXA1 and AP1 appears to have a dramatic effect on FOXA1 activity. In the presence of AP1 sites, FOXA1 displays anticooperative binding dynamics where occupancy decreases at the highest levels of FOXA1 expression. We speculate that at these sites monomers of FOXA1 interact with AP1 to potentiate binding, whereas dimers of FOXA1 cannot cobind with AP1. In this model, high concentrations of FOXA1 favor its dimeric form which accounts for the loss of binding at these sites when FOXA1 is expressed at high levels. Regardless of the mechanism underlying anticooperative behavior, our results show that pioneer activity can be modified by the interactions a TF makes inside cells. Thus, pioneer activity is contingent on many properties of a TF including its levels, its intrinsic affinity for its motif, the motif content at its targets, and the different interactions it makes with other proteins when bound at different locations. Given these contingencies we suggest that most TFs will display some degree of pioneer activity and that the Δdox_50_, or a related metric, will be a useful metric to quantify it.

## Materials and Methods

### Cell lines

We grew K562 cells (ATCC CCL-243, Manassas, VA) in Iscove’s Modified Dulbecco Serum supplemented with 10% fetal bovine serum, 1% penicillin-streptomycin and 1% non-essential amino acids. We used these cell types to generate clonal FOXA1 and HNF4A lines, as described below. For each of our functional assays, we split each line into replicate flasks, treated with doxycycline (dox) (Sigma #D9891-1G), and then waited 24 hours to extract RNA or nuclei. We used doses of 0.005μg/ml, 0.05μg/ml, 0.1μg/ml, 0.25μg/ml, 0.5μg/ml, and 5μg/ml for our dox_50_ experiments.

### Cloning, production, and infection of viral vectors

We used FOXA1 and HNF4A K562 clonal lines and lentiviral vectors carrying inducible FOXA1 and HNF4A ORFs as described previously (Hansen, Loell, and Cohen 2022).

### Sequencing library preparations and analysis

We prepared sequencing libraries and analyzed the two replicates of CUT&Tag as described previously (Hansen, Loell, and Cohen 2022). In our previous work we already used ATAC-seq to measure the uninduced (-dox) accessibility in the FOXA1 and HNF4A K562 lines (Hansen, Loell, and Cohen 2022). Because we used the same clones to perform these experiments, we re-used these data as uninduced accessibility. We also had already sequenced CUT&Tag libraries for the 0.5ug/ml and 0.05ug/ml doxycycline induction levels and re-used these data as well.

### Binding curve analysis

We first established a set of all possible binding sites for each TF by creating a list of binding sites in the sample with the highest dox induction concentration (5μg/ml). We subset this list into those accessible binding sites (called accessible peak in the -dox uninduced condition) and inaccessible binding sites (absence of called accessible peak). Then we used the multiBigwigSummary from the deepTools suite (Ramírez et al. 2016) to count the normalized read intensity at each peak from each induction level. We normalized each induction level to the read intensity at the highest induction level in order to convert read intensity into fraction bound.

With these data, we fit a binding curve using SciPy curvefit (Virtanen et al. 2020) to the equation (Equation 1) where dox_50_ is unknown and represents a binding affinity parameter similar to K_d_ and where [dox] is the concentration of dox used to induce TF expression. When we plotted examples of randomly selected genomic sites and examined the binding curves, we noticed that at some sites, binding peaked (fraction bound >= 1) prior to the highest concentration. In these cases, the fit line estimated a negative dox_50_. For this reason, we filtered out any site that peaked prior to the sample with the highest dox concentration. We also estimated dox_50_ distributions without this filtering step and found similar distributions. (Fig. S2).

In order to quantify the early peak, or “anti-cooperative” behavior that we observed, we classified a binding site as exhibiting a“saturation binding” modality if only the highest dox concentration had a fraction bound of 1, and then each subsequent lower concentration had a lower fraction bound. We classified a binding site as exhibiting an “anti-cooperative” modality if the site peaked at either the third (0.1μg/ml) or fourth (0.25μg/ml) dox concentrations and then declined in each direction.

We calculated reproducibility in three ways. We first showed that the binding signal was reproducible by plotting the RPKM signal from each replicate for each of the concentrations at all of the binding sites collected as described above. We then showed that the lines fit similarly between replicates by both replicates’ binding signal and fit binding curves at many different randomly chosen genomic sites and showing that the lines look similar. And finally we showed that the distributions of dox_50_s from each replicate were highly overlapping. After showing these, we averaged the dox_50_ from each replicate at each site and used the average value moving forward.

### Motif analysis

To discover or count motifs in binding sites, we extracted the sequence from each CUT&Tag binding peak and then used XSTREME (Grant and Bailey 2021) for de novo motif discovery and FIMO (Grant, Bailey, and Noble 2011) for specific motif occurrence counting. We used 1e-3 as a p-value threshold and JASPAR (Fornes et al. 2020) PWMs for FOXA1 (MA0148.1), HNF4A (MA0114.2), and AP-1 (MA1141.1). We used these motif counts to subset the FOXA1/HNF4A accessible/inaccessible peaks into those with less than 2 motifs, more than 2 but less than 4, or 4 or more, and then re-ran the analysis (Fig. S4).

### Chromatin modifications analysis and modeling

We used previously published datasets of histone ChIP-seq (Zhang et al. 2020) to identify patterns of H3K27Ac, H3K4me1, H3K9me3, and H3K27me3 marks. We used BEDTools (Quinlan and Hall 2010) to overlap FOXA1 or HNF4A’s binding sites with these marks. We then used python’s statsmodels to run ANOVA analyses on ordinary least squares linear regressions. Each reported variance is the parameter’s sum of squares contribution divided by the total sum of squares.

## Data Availability

All genomic sequencing data have been submitted to Gene Expression Omnibus (GEO). We are awaiting review and assignment of an accession number.

## Acknowledgements

We thank members of the Cohen Lab for reading and critiquing the manuscript and for helpful discussion; Jessica Hoisington-Lopez and MariaLynn Crosby in the DNA Sequencing Innovation Lab for assistance with high-throughput sequencing; the Genome Engineering and iPSC Center for allowing us to use their Sony Flow Cytometer for cell sorting; and Mingjie Li in the Hope Center Viral Vectors Core for assistance with producing lentiviral expression vectors. This work was supported by grants from the National Institutes of Health: R01GM092910 (Dr. Barak Cohen), T32HG000045 (Dr. Michael Brent, Washington University in St. Louis Genome Analysis Training Program), and T32GM007200 (Dr. Wayne Yokoyama, Washington University in St. Louis Medical Scientist Training Program).

## Author Contributions

J.L.H. and B.A.C. designed the overall project. J.L.H. conducted experiments. J.L.H conducted analysis. J.L.H. and B.A.C. wrote the manuscript.

## Competing Interests

The authors declare no competing interests.

## Notes

### Competing Interest Statement

The authors have declared no competing interest.

## References

Biddie, Simon C., Sam John, Pete J. Sabo, Robert E. Thurman, Thomas A. Johnson, R. Louis Schiltz, Tina B. Miranda, et al. 2011. “Transcription Factor AP1 Potentiates Chromatin Accessibility and Glucocorticoid Receptor Binding.” Molecular Cell 43 (1): 145–55.

Biddy, Brent A., Wenjun Kong, Kenji Kamimoto, Chuner Guo, Sarah E. Waye, Tao Sun, and Samantha A. Morris. 2018. “Single-Cell Mapping of Lineage and Identity in Direct Reprogramming.” Nature 564 (7735): 219–24.

Buenrostro, Jason D., Beijing Wu, Howard Y. Chang, and William J. Greenleaf. 2015. “ATAC-Seq: A Method for Assaying Chromatin Accessibility Genome-Wide.” Current Protocols in Molecular Biology / Edited by Frederick M. Ausubel…[et Al.] 109 (January): 21.29.1–9.

Cirillo, Lisa Ann, Frank Robert Lin, Isabel Cuesta, Dara Friedman, Michal Jarnik, and Kenneth S. Zaret. 2002. “Opening of Compacted Chromatin by Early Developmental Transcription Factors HNF3 (FoxA) and GATA-4.” Molecular Cell 9 (2): 279–89.

Creyghton, Menno P., Albert W. Cheng, G. Grant Welstead, Tristan Kooistra, Bryce W. Carey, Eveline J. Steine, Jacob Hanna, et al. 2010. “Histone H3K27ac Separates Active from Poised Enhancers and Predicts Developmental State.” Proceedings of the National Academy of Sciences of the United States of America 107 (50): 21931–36.

Fornes, Oriol, Jaime A. Castro-Mondragon, Aziz Khan, Robin van der Lee, Xi Zhang, Phillip A. Richmond, Bhavi P. Modi, et al. 2020. “JASPAR 2020: Update of the Open-Access Database of Transcription Factor Binding Profiles.” Nucleic Acids Research 48 (D1): D87–92.

Garcia, Meilin Fernandez, Cedric D. Moore, Katharine N. Schulz, Oscar Alberto, Greg Donague, Melissa M. Harrison, Heng Zhu, and Kenneth S. Zaret. 2019. “Structural Features of Transcription Factors Associating with Nucleosome Binding.” Molecular Cell. https://doi.org/10.1016/j.molcel.2019.06.009.

Grant, Charles E., and Timothy L. Bailey. 2021. “XSTREME: Comprehensive Motif Analysis of Biological Sequence Datasets.” bioRxiv. https://doi.org/10.1101/2021.09.02.458722.

Grant, Charles E., Timothy L. Bailey, and William Stafford Noble. 2011. “FIMO: Scanning for Occurrences of a given Motif.” Bioinformatics 27 (7): 1017–18.

Hansen, Jeffrey L., Kaiser J. Loell, and Barak A. Cohen. 2022. “The Pioneer Factor Hypothesis Is Not Necessary to Explain Ectopic Liver Gene Activation.” eLife 11 (January). https://doi.org/10.7554/eLife.73358.

Heintzman, Nathaniel D., Rhona K. Stuart, Gary Hon, Yutao Fu, Christina W. Ching, R. David Hawkins, Leah O. Barrera, et al. 2007. “Distinct and Predictive Chromatin Signatures of Transcriptional Promoters and Enhancers in the Human Genome.” Nature Genetics 39 (3): 311–18.

Horisawa, Kenichi, Miyako Udono, Kazuko Ueno, Yasuyuki Ohkawa, Masao Nagasaki, Sayaka Sekiya, and Atsushi Suzuki. 2020. “The Dynamics of Transcriptional Activation by Hepatic Reprogramming Factors.” Molecular Cell 79 (4): 660–76.e8.

Iwafuchi-Doi, Makiko, and Kenneth S. Zaret. 2014. “Pioneer Transcription Factors in Cell Reprogramming.” Genes & Development 28 (24): 2679–92.

Jiang, G., U. Lee, and F. M. Sladek. 1997. “Proposed Mechanism for the Stabilization of Nuclear Receptor DNA Binding via Protein Dimerization.” Molecular and Cellular Biology 17 (11): 6546–54.

Kaya-Okur, Hatice S., Steven J. Wu, Christine A. Codomo, Erica S. Pledger, Terri D. Bryson, Jorja G. Henikoff, Kami Ahmad, and Steven Henikoff. 2019. “CUT&Tag for Efficient Epigenomic Profiling of Small Samples and Single Cells.” Nature Communications 10 (1): 1930.

Man, T. K., and G. D. Stormo. 2001. “Non-Independence of Mnt Repressor-Operator Interaction Determined by a New Quantitative Multiple Fluorescence Relative Affinity (QuMFRA) Assay.” Nucleic Acids Research 29 (12): 2471–78.

Mayran, Alexandre, Konstantin Khetchoumian, Fadi Hariri, Tomi Pastinen, Yves Gauthier, Aurelio Balsalobre, and Jacques Drouin. 2018. “Pioneer Factor Pax7 Deploys a Stable Enhancer Repertoire for Specification of Cell Fate.” Nature Genetics 50 (2): 259–69.

Miller, Joanna A., and Jonathan Widom. 2003. “Collaborative Competition Mechanism for Gene Activation in Vivo.” Molecular and Cellular Biology 23 (5): 1623–32.

Quinlan, Aaron R., and Ira M. Hall. 2010. “BEDTools: A Flexible Suite of Utilities for Comparing Genomic Features.” Bioinformatics 26 (6): 841–42.

Ramírez, Fidel, Devon P. Ryan, Björn Grüning, Vivek Bhardwaj, Fabian Kilpert, Andreas S. Richter, Steffen Heyne, Friederike Dündar, and Thomas Manke. 2016. “deepTools2: A next Generation Web Server for Deep-Sequencing Data Analysis.” Nucleic Acids Research 44 (W1): W160–65.

Rufibach, Laura E., Stephen A. Duncan, Michele Battle, and Samir S. Deeb. 2006. “Transcriptional Regulation of the Human Hepatic Lipase (LIPC) Gene Promoter.” Journal of Lipid Research 47 (7): 1463–77.

Sekiya, Sayaka, and Atsushi Suzuki. 2011. “Direct Conversion of Mouse Fibroblasts to Hepatocyte-like Cells by Defined Factors.” Nature 475 (7356): 390–93.

Soufi, Abdenour, Meilin Fernandez Garcia, Artur Jaroszewicz, Nebiyu Osman, Matteo Pellegrini, and Kenneth S. Zaret. 2015. “Pioneer Transcription Factors Target Partial DNA Motifs on Nucleosomes to Initiate Reprogramming.” Cell 161 (3): 555–68.

Swinstead, Erin E., Tina B. Miranda, Ville Paakinaho, Songjoon Baek, Ido Goldstein, Mary Hawkins, Tatiana S. Karpova, et al. 2016. “Steroid Receptors Reprogram FoxA1 Occupancy through Dynamic Chromatin Transitions.” Cell 165 (3): 593–605.

Virtanen, Pauli, Ralf Gommers, Travis E. Oliphant, Matt Haberland, Tyler Reddy, David Cournapeau, Evgeni Burovski, et al. 2020. “SciPy 1.0: Fundamental Algorithms for Scientific Computing in Python.” Nature Methods 17 (3): 261–72.

Yan, Chao, Hengye Chen, and Lu Bai. 2018. “Systematic Study of Nucleosome-Displacing Factors in Budding Yeast.” Molecular Cell 71 (2): 294–305.e4.

Yu, Xinyang, and Michael J. Buck. 2019. “Defining TP53 Pioneering Capabilities with Competitive Nucleosome Binding Assays.” Genome Research 29 (1): 107–15.

Zhang, Jing, Donghoon Lee, Vineet Dhiman, Peng Jiang, Jie Xu, Patrick McGillivray, Hongbo Yang, et al. 2020. “An Integrative ENCODE Resource for Cancer Genomics.” Nature Communications 11 (1): 3696.

Zhu, Fangjie, Lucas Farnung, Eevi Kaasinen, Biswajyoti Sahu, Yimeng Yin, Bei Wei, Svetlana O. Dodonova, et al. 2018. “The Interaction Landscape between Transcription Factors and the Nucleosome.” Nature 562 (7725): 76–81.

