## Supplemental Information for "A quantitative metric of pioneer activity reveals that HNF4A has stronger in vivo pioneer activity than FOXA1"

### Supplementary Information

### Supplementary Figures

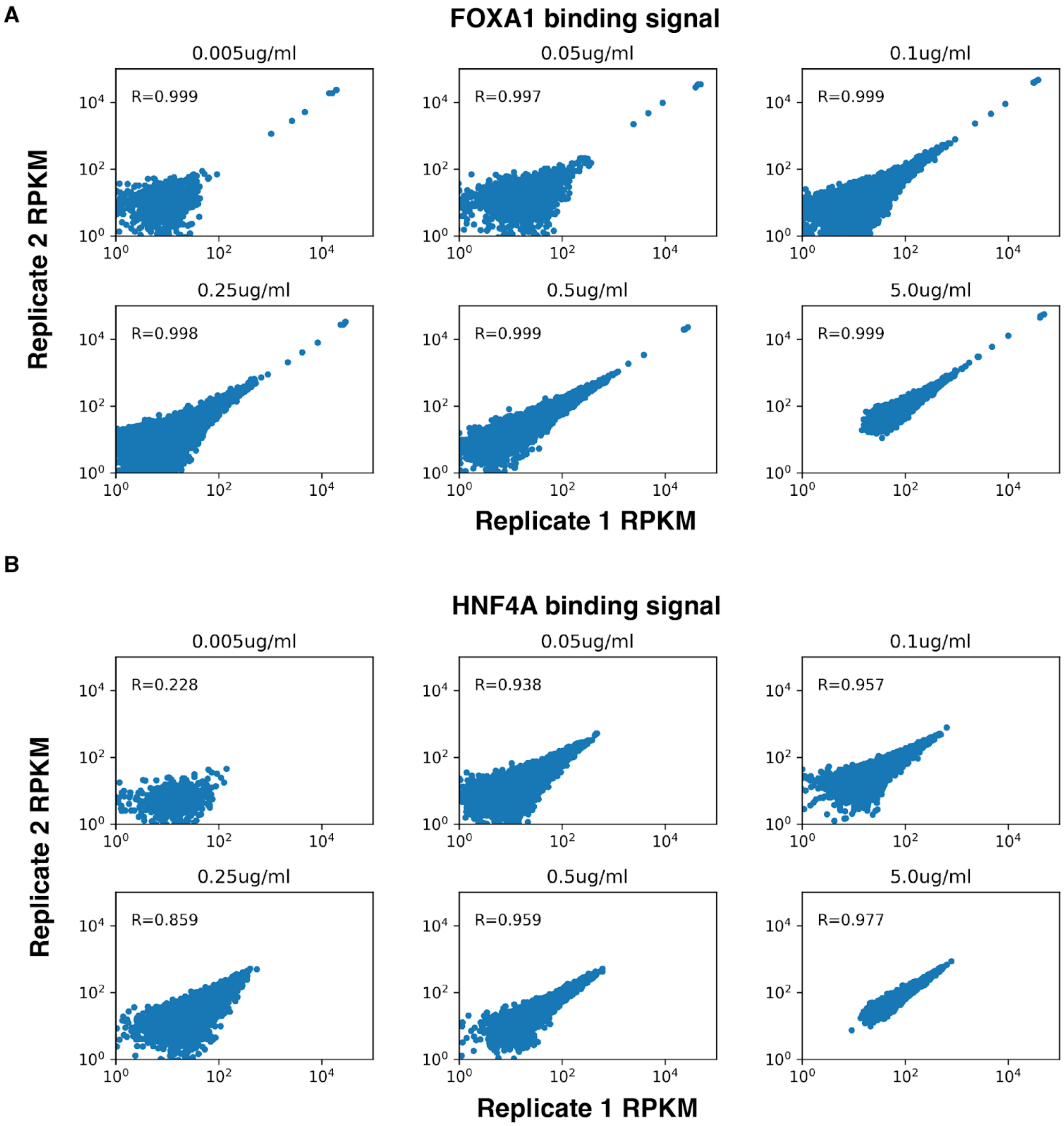


**Figure S1. Reproducibility of binding signal.** RPKM signal from each replicate of CUT&Tag data across each TF across each dox induction concentration. Pearson's R correlation displayed on each graph.

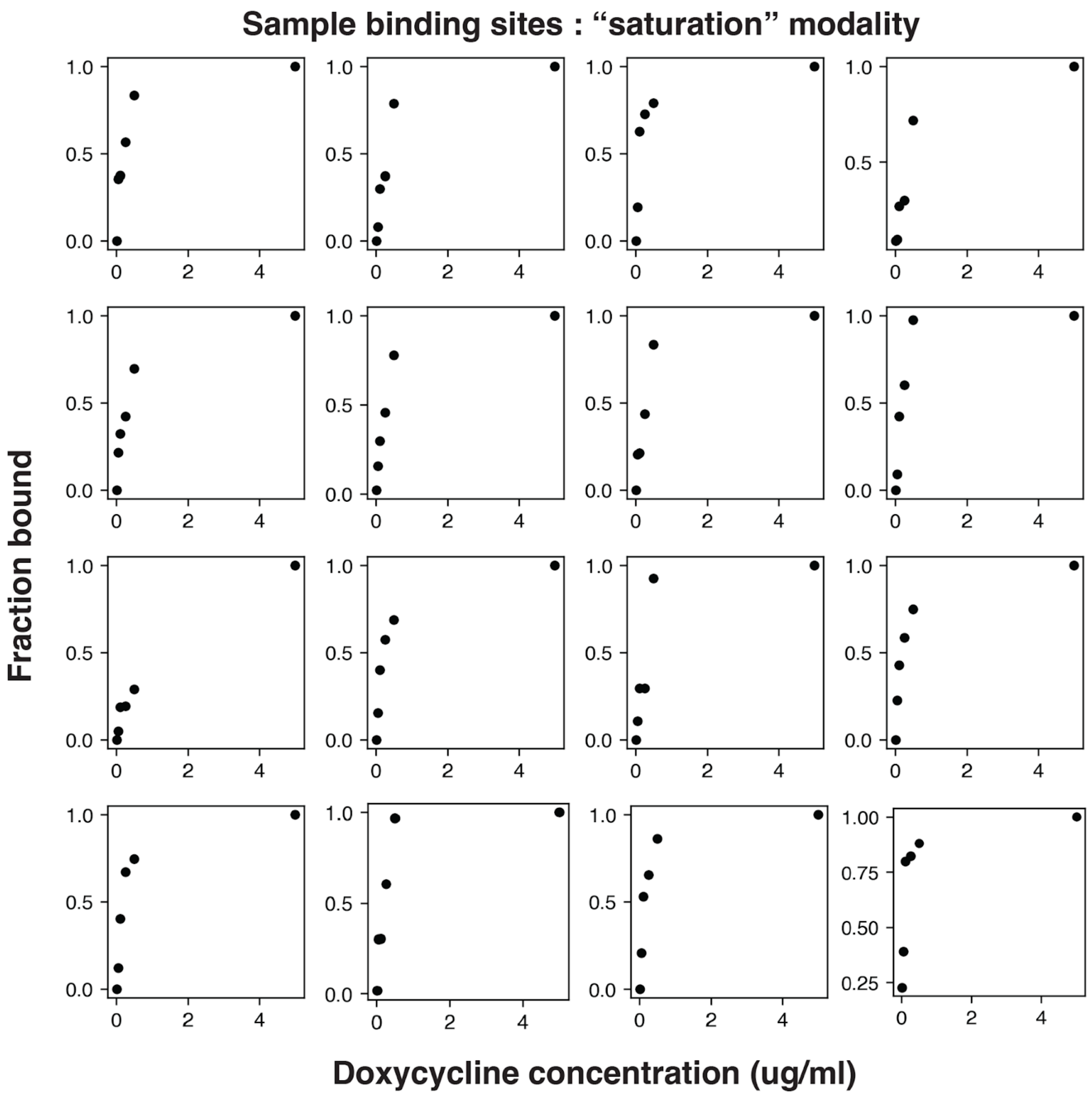


**Figure S2. Common saturation behavior binding pattern.** 16 examples from different genomic sites showing saturating binding signal as dox induction increases. Signal is first read normalized (RPKM) and then normalized to the signal at the highest concentration. These sites were sampled from FOXA1 accessible binding sites, but are common across inaccessible and HNF4A binding sites as well.

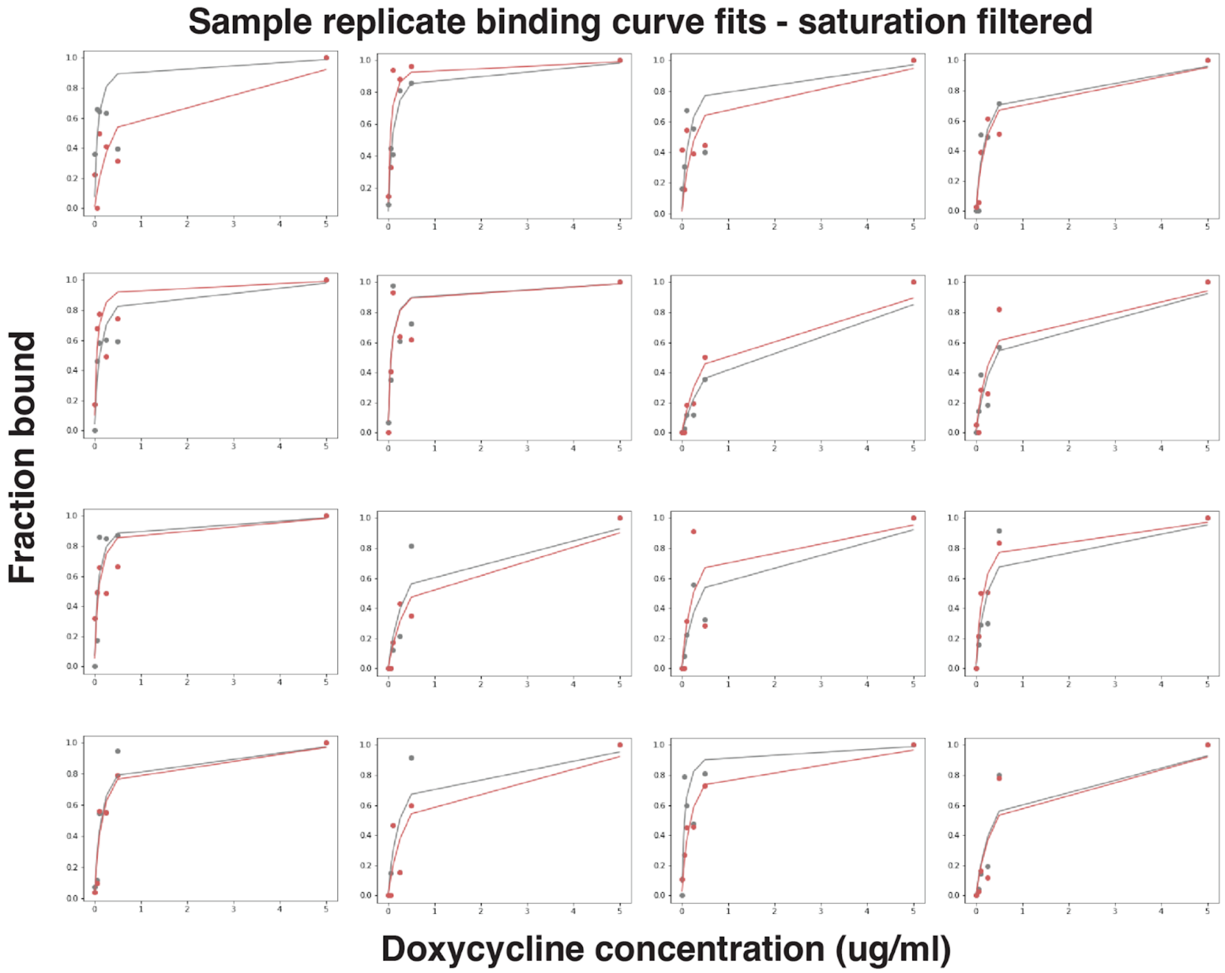


**Figure S3. Sample of replicate fit binding curves.** RPKM binding signal and fitted lines for each CUT&Tag replicate at 16 representative genomic loci

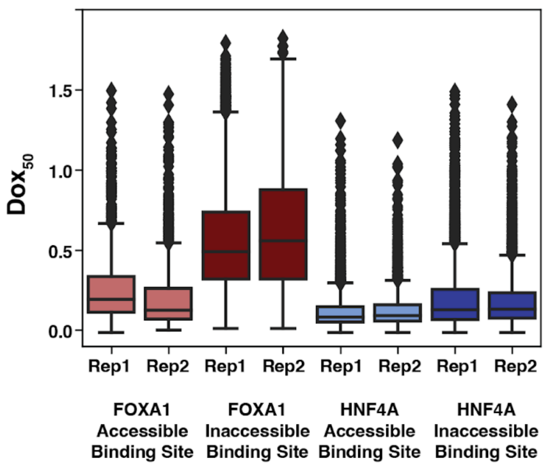

**Figure S4. Replicate $dox_{50}$ distributions.** $Dox_{50}$ distributions extracted from fitted lines from each CUT&Tag replicate across each TF and each type of chromatin.

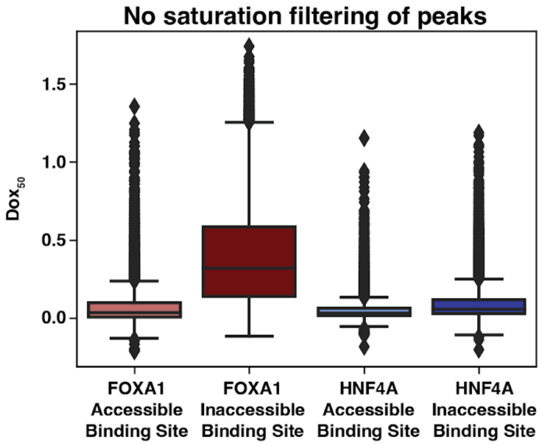


**Figure S5. $Dox_{50}$ distributions without filtering out early saturation peaks.** Dox50 distributions from all of the FOXA1 accessible binding sites (*n* = 10,118), FOXA1 inaccessible binding sites (*n* = 17,644), HNF4A accessible binding sites (*n* = 16,137), and HNF4A inaccessible binding sites (*n* = 16,507), without filtering out those peaks where binding signal peaked prior to the 5ug/ml dox sample.

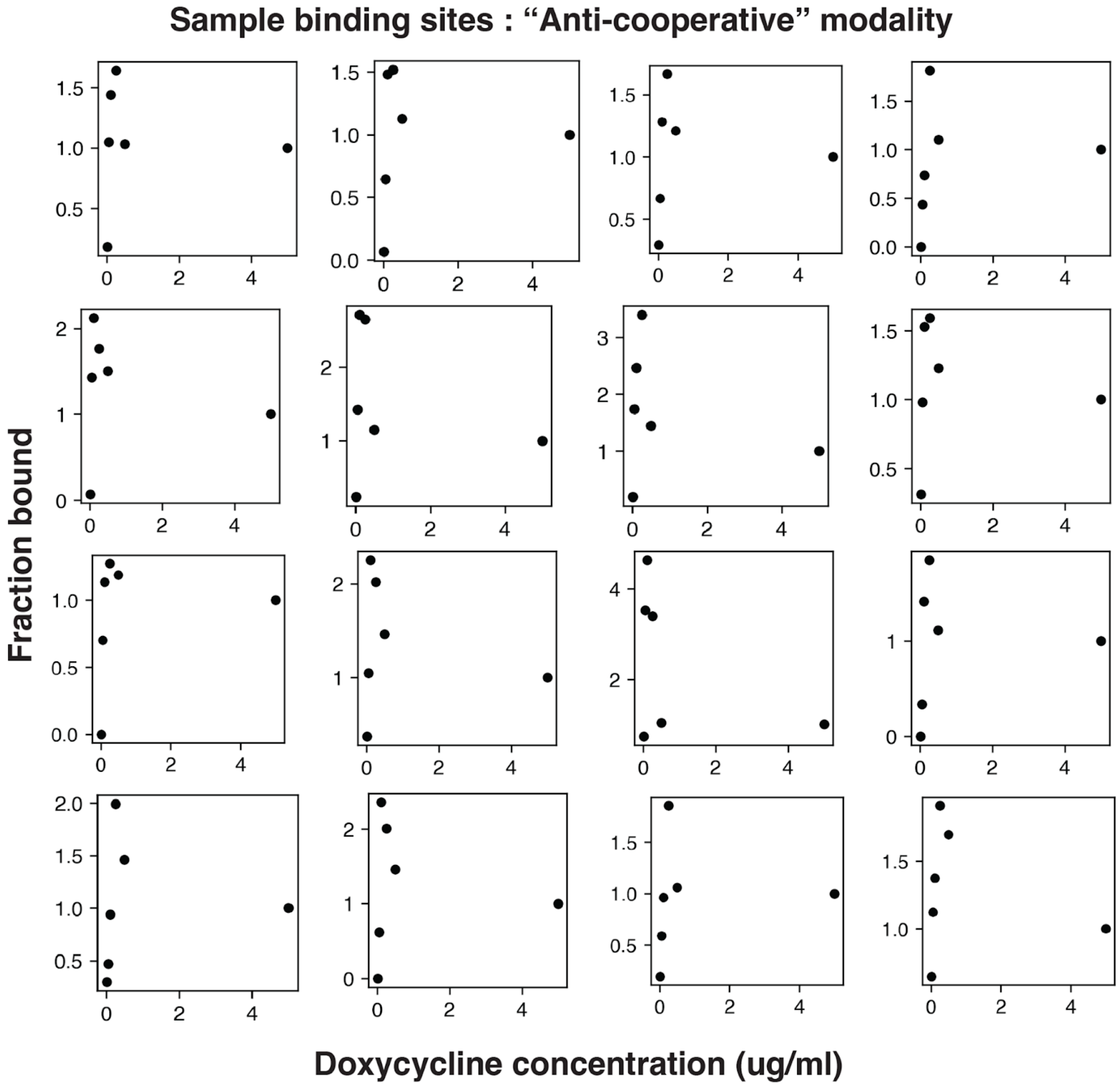


**Figure S6. Common "anti-cooperative" binding pattern.** 16 examples from different genomic sites showing a pattern of increasing and then decreasing binding signal as dox induction increases. Signal is first read normalized (RPKM) and then normalized to the signal at the highest concentration. These sites were sampled from FOXA1 accessible binding sites.
